# Genome wide association study of Arabidopsis seed mucilage layers at a regional scale

**DOI:** 10.1101/2023.10.21.563161

**Authors:** Sébastien Viudes, Rémy Zamar, Vincent Burlat, Fabrice Roux, Christophe Dunand

## Abstract

The myxospermous species *Arabidopsis thaliana* extrudes a polysaccharidic mucilage from the seed coat epidermis during imbibition. The whole seed mucilage can be divided into a seed-adherent layer and a fully soluble layer, both layers presenting natural genetic variations. The adherent mucilage is variable in size and composition, while the soluble mucilage is variable in composition and physical properties. Studies reporting both the genetic architecture and the putative selective agents acting on this natural genetic variation are scarce. In this study, we set up a Genome Wide Association study (GWAS) based on 424 natural accessions collected from 166 natural populations of *A. thaliana* located south-west of France and previously characterized for a very important number of abiotic and biotic factors. We identified an extensive genetic variation for both mucilage layers. The adherent mucilage was mainly related to precipitation and temperature whereas the non-adherent mucilage was unrelated to any environmental factors. By combining a hierarchical Bayesian model with a local score approach, we identified 55 and 28 candidate genes, corresponding to 26 and 10 QTLs for the adherent and non-adherent mucilages, respectively. Putative or characterized function and expression data available in the literature were used to filter the candidate genes. Only one gene among our set of candidate genes was already described as a seed mucilage actor, leaving a large set of new candidates putatively implicated inseed mucilage synthesis or release. The present study lay out foundation to understand the influence of regional ecological factors acting on seed mucilage in *A. thaliana*.

## 1 Introduction

Seed mucilage is a polysaccharidic hydrogel extruded around seeds upon imbibition by the outermost epidermal cell layer of the seed coat in myxospermous species (Phan and Burton, 2018). Seed mucilage can help plant development from germination until seedling establishment in biotic and abiotic stress conditions (Viudes et al., 2020). Depending on plant species, the interaction between seed mucilage and biotic stresses can be direct, e.g. escaping ant predation by sticking to soil (Pan et al., 2021) or facilitating passage through the digestive tract of birds (Kreitschitz et al., 2021), or indirect, e.g. mobilizing soil bacteria that limit the development of plant pathogen fungi, which lead to reduced disease symptoms and a better plant development (Meschke and Schrempf, 2010). Concerning abiotic stresses, seed mucilage is known to influence or to rely on water around the seed by regulating the incoming flow, and by changing soil water retention capacity (Deng et al., 2014; Saez-Aguayo et al., 2014; Teixeira et al., 2020). The ecological function of seed mucilage was historicaly studied in several plant species. In contrast, the genetics and development of seed mucilage was nearly exclusively studied in the model plant *Arabidopsis thaliana*.

In *A. thaliana*, around one hundred genes are participating to mucilage secretory cell formation and mucilage biosynthesis and release (Viudes et al., 2021). A large majority of these genes have been characterized through classical forward and reverse genetic studies. First of these were major regulators or main actors of mucilage establishment that lead to drastic phenotypes when mutated (Western, 2001). Progressively, the newly discovered genes had more subtle roles for myxospermy, displaying more discrete phenotypes such as specific polysaccharide proportions or decorations when mutated.

Evidence of natural diversity for seed mucilage was first shown with the characterisation of an *A. thaliana* natural genotype, originated from Tajikistan, which is not able to extrude mucilage (Macquet et al., 2007). Among 22 *A. thaliana* populations originating from central Asia and Scandinavia, seven were reported to be fully defective for mucilage release (Saez-Aguayo et al., 2014). Total seed mucilage can have great polysaccharide composition changes among natural populations, regarding mannose, galactose, galacturonic acid and rhamnose monosaccharide contents (Voiniciuc et al., 2016). The *A. thaliana* mucilage has two distinct layers, the inner layer is highly adherent to the seed thanks to cellulosic microfibrils, and the outer layer is fully soluble (Tsai et al., 2017). The volume of adherent mucilage was shown to be also variable among natural populations probably linked to the observed cellulose crystalline structure changes (Voiniciuc et al., 2016). The non-adherent mucilage exhibits also variable compositions and physical properties (Poulain et al., 2019). Interestingly, the intra-species loss of myxospermy can occur several times due to independent genetic mutations (Saez-Aguayo et al., 2014). Two independent Genome-Wide Association Studies (GWAS), based on *A. thaliana* adherent and non-adherent seed mucilage phenotyping, highlighted 21 and 11 significant SNPs, respectively, supporting the polygenic architecture of myxospermy (Fabrissin et al., 2019; Voiniciuc et al., 2016). Due to the lack of availability of accurate environmental data for many *A. thaliana* natural populations, natural genetic variation in mucilage characteristics has so far never been associated with environmental data through a statistical approach.

Here, we aimed at using GWAS to further investigate the genetics underlying natural genetic variation of *A. thaliana* myxospermy. Our hypothesis was that this approach without *a priori* will highlight new candidate genes that were not yet linked to seed mucilage. The quantitative and independent phenotyping of both mucilage layers should increase the efficiency of GWAS and also provide the opportunity to study if a correlation exists between the two layers of mucilage. In addition, the availability of climatic conditions and soil composition for the studied populations created a unique opportunity to address the potential ecological role of myxospermy in *A. thaliana*.

## 2 Material and methods

### 2.1 Plant material

Our study was based on 168 natural populations of *A. thaliana* identified in spring 2014 and located in the Midi-Pyrénées region (south-west of France) (Bartoli et al., 2018; Frachon et al., 2018). The average distance between these populations is ∼100 km. These populations have been characterized for six non-correlated variables retrieved from the ClimateEU database (Frachon et al., 2018), 13 chemical and 1 physical soil properties with soil samples collected *in situ* in late autumn 2015 and late winter 2015(Frachon et al., 2019), leaf and root bacterial microbiota and pathobiota in late autumn 2015 and late winter 2015(Bartoli et al., 2018), and plant communities in late spring 2015 (Frachon et al., 2019). In addition, based on a Pool-Seq approach with an average of 15.3 plants per population, the 168 populations have been whole-genome sequenced with the Illumina technology, leading to the identification of 1,638,649 SNPs (Frachon et al., 2018). For the purpose of this study, we used 166 out of the 168 initial populations, each being composed of one to three accessions, thereby bringing the total of phenotyped seed lots to 424. Differences in the maternal effects among the 424 seed lots were reduced by growing one plant of each accession for one generation with the following steps: (i) for each accession, several seeds were sown in one 7 x 7 x 6 cm plastic pot (Soparco®, Condé-sur-Huisne, France) filled with damp standard culture soil (PROVEEN MOTTE 20, Soprimex®, L’Isle-d’Abeau, France) on November 1^st^ 2016, (ii) seeds were stratified at 4°C for four days in order to break primary dormancy and promote germination, (iii) pots were transferred to greenhouse conditions (22°C, 16 h photoperiod) on November 4^th^ 2016, (iv) seedlings were thinned to one on November 25^th^ 2016, (v) pots were placed on a field station at the INRAE campus of Castanet-Tolosan (France) on December 5^th^ 2021 in order to promote flowering through natural vernalization, (vi) at the onset of flowering, plants were transferred in a greenhouse mimicking the outdoor conditions (no additional light and heating) but protecting plants from rainfall, and (vii) plants were separated from each other by aratubes in order to avoid cross-pollination among accessions. All seed lots were harvested during a 3-week period from late April to early May 2017 and were then stored at 4°C until the set-up of the phenotyping experiments.

### 2.2 Phenotyping experiments

Given the high number of accessions, the experiments were performed sequentially with several experimental batches distributed along a few months and randomly selected. The number of accessions in each batch was dependent on the phenotypic traits. In order to consider micro-environmental variations among the experimental batches, the same Columbia (Col-0) seed lot was used as an internal control that was systematically included in all experimental batches in order to normalize the results.

#### 2.2.1 Adherent mucilage and seed area phenotyping

A total of 44 experimental batches, including Col-0 as a control and 11 different accessions, were set up during over 6 months. Because the adherent mucilage is strongly attached to the seed, we used a previously described protocol that allows comparing adherent mucilage area among accessions in the most standardized manner (Francoz et al., 2019). Briefly, to remove the non-adherent mucilage, 50 to 100 seeds per accession were vigorously shaken at 250 rpm in 1.6 ml of 0.01 M Tris-HCl pH 7.5 contained in a 2 ml tube. The remaining adherent layer of mucilage was then stained with a 0.02 % (w/v) ruthenium red solution in Tris-HCl. The seeds were rinsed in 0.01 M Tris-HCl pH 7.5, transferred in 24-well microplates containing 0.01 M Tris-HCl pH 7.5 and imaged using an Epson perfection V100 photo scanner at 6400 dpi resolution. The measurements of seed area including or not the adherent mucilage was realized with imageJ using a previously described macro (Francoz et al., 2019). The adherent mucilage area was calculated for each seed by subtracting the seed area alone from the seed area with the adherent mucilage. For each experimental batch, raw data for adherent mucilage area and seed area of each accession were standardized with the corresponding means of adherent mucilage and seed areas of Col-0.

#### 2.2.2 Non-adherent mucilage area phenotyping

A total of 98 batches of experiments including Col-0 as a control and three or six different accessions, were set up over 6 months. The non-adherent mucilage is totally soluble in water and quickly released upon seed imbibition. To estimate its area, we used the MuSeeQ protocol (Miart et al., 2018). Briefly, 5 to 10 dry seeds by accession were individually and synchronously disposed on perfectly flat 0.6 % (w/v) agarose media containing 0.00004 % (w/v) of toluidine blue O polychromatic stain (Sigma-Aldrich, Saint-Quentin-Fallavier, France). Upon the contact with the aqueous media, the non-adherent mucilage is released, spreading around the seed on the flat surface, and is simultaneously stained in pink contrasting with the blue colour of the media. Top-view pictures were taken 24 h after seed deposition with a camera (Canon DS 126271). The non-adherent mucilage area was measured for each seed using imageJ with a previously described macro (Miart et al., 2018). For each experimental batch, raw data for non-adherent mucilage area of each accession were standardized with the corresponding mean of Col-0 non-adherent mucilage area.

### 2.3 Statistical analyses

#### 2.3.1 Natural genetic variation

To estimate the extent of natural genetic variation for each mucilage trait, we run the following mixed models using the PROC MIXED procedure in SAS v9.4 (SAS Institute Inc., Cary, North Carolina, USA):

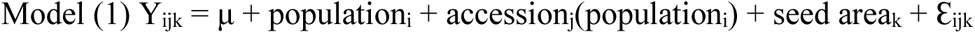

Where ‘Y’ is the mucilage area of seeds, ‘μ’ is the overall mean of mucilage area, ‘population’ accounts for genetic differences among the 166 populations, ‘accession(population)’ accounts for the mean genetic difference among accessions within population, ‘seed area’ is a covariate to control for the developmental effect of seed size on adherent mucilage area, and ‘Ɛ’ is the residual term. The factor ‘population’ was treated as a fixed effect, whereas the factor ‘accession’ was treated as a random effect. Following Brachi et al. (2013), this model random term was tested with a likelihood ratio test (LRT) with and without this effect.

For both phenotypic traits, we estimated the percentage of variance observed among and within populations by running Model 1 with the VARCOMP procedure in SAS v9.4 (SAS Institute Inc., Cary, North Carolina, USA). In addition, we estimated genotypic values of the 166 natural populations by calculating least-square means (LSmeans) of each population after removing the ‘accession(population)’ term from Model 1.

Spearman’s correlation coefficients (Spearman’s *rho*) were calculated to estimate relationships between genetic differences among populations (i.e. genotypic values) and abiotic ecological variables. Resulting *p*-values were corrected with an FDR (False Discovery Rate) correction.

#### 2.3.2 GWA mapping with local score analysis

To detect QTLs associated with natural genetic variation of mucilage area related traits, we combined a GWA mapping approach with a local score approach. This combination of approaches already allowed the successful detection and cloning in *A. thaliana* of five QTLs in response to bacterial pathogens (Aoun et al., 2020; Demirjian et al., 2022; Demirjian et al., 2023). First, we retrieved for each of the 166 populations the standardized allele frequencies for all the SNPs (see ‘Plant material’ subsection). These standardized allele frequencies result from raw allele frequencies after accounting for the scaled covariance of population allele frequencies, thereby making the phenotype-genotype associations robust to complex demographic histories and allowing decreasing drastically the rate of false positives (Frachon et al., 2019, 2018). Second, for each phenotypic trait, we estimated phenotype-genotype associations across the genome by calculating Spearman correlation coefficients between standardized allele frequencies of each SNP and genotypic values of the 166 populations, using the *cor.test* function implemented under the *R* environment. Third, we implemented a local score approach on the set of *p*-values associated with the Spearman’s *rho* values. The local score allows accumulating the statistical signals from contiguous genetic markers in order to detect significant genomic regions associated with phenotypic natural variation (Fariello et al., 2017). Through the *p*-values in a given QTL region, the association signal will cumulate locally due to linkage disequilibrium between SNPs, which will then increase the local score (Bonhomme et al., 2019). The tuning parameter ξ was fixed at 2 and significant SNP-phenotype associations were identified by estimating a chromosome-wide significance threshold for each chromosome (Bonhomme et al., 2019).

Based on a custom script described in Libourel et al. (2021), we retrieved all candidate genes underlying QTLs by selecting all genes inside the QTL regions and also the first gene upstream and the first gene downstream of these QTL regions. The TAIR 10 database (https://www.arabidopsis.org/) was used as a reference.

### 2.4 Transcriptomic profiling of candidate genes

The expression profiles of the candidate genes were extracted from *A. thaliana* tissue-specific seed development transcriptomic data (Belmonte et al., 2013). Whole transcriptomic data set used to decipher the seed coat specific expression of the candidate genes are available in Supplementary Table S2.

## 3 Results

### 3.1 Large genetic variation among and within natural populations for seed mucilage

The sizes of adherent and non-adherent mucilage layers were measured for each accession belonging to the 166 populations (Fig. 1). Although the two different mucilages exist in 3D, their quantifications come from 2D images. Mucilage values are therefore expressed in surface unit. For adherent mucilage, measured values correspond to orthogonal projections, which are coherent with the reality since the seeds systematically lay down longitudinally in solution due to their oblong shape. Non-adherent mucilage spreads on a flat surface and is consequently already a 2D object before the imaging.

**Fig. 1.**
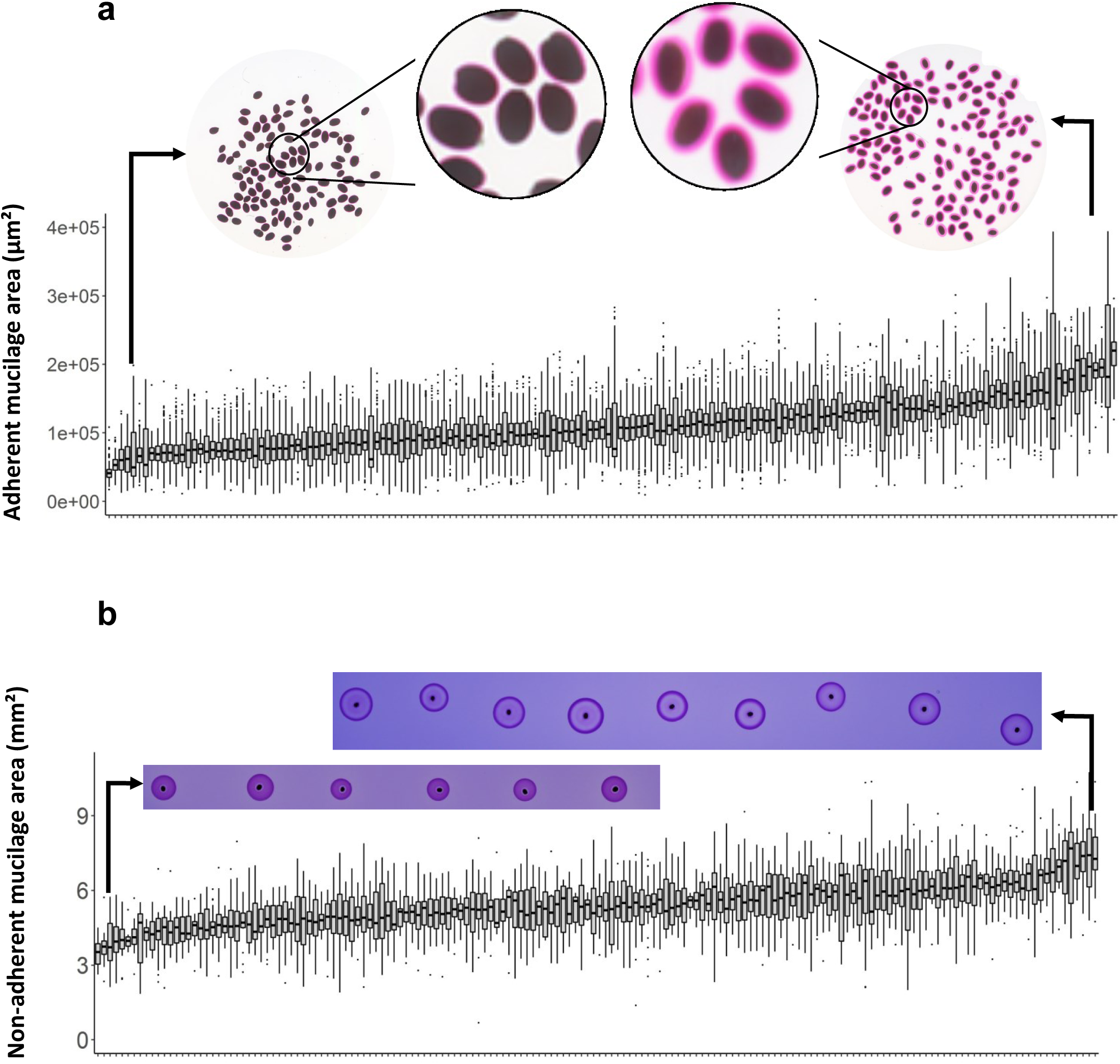
Measurement of adherent **(a)** and non-adherent mucilage **(b)** of each accession belonging to the 166 natural populations and source picture examples illustrating the extreme area values. Each individual boxplot represents the measured area distribution (µm²) of the adherent mucilage within a population. Each population contains one to three accession. For each accession 50 to 100 seeds or 5 to 10 seeds are phenotyped for adherent and non-adherent mucilage respectively. Boxplots are ordered by their mean value. The images correspond to the extreme values to highlight the existing contrast. Measurement were realized on such images after vigorous shaking and ruthenium red staining or 24 h after dry seed deposition on a Toluidine blue agarose media for adherent and non-adherent mucilage respectively.

For both mucilage layers, a large phenotypic variation was observed among the 166 populations (Fig. 1). The ratio of the adherent-mucilage area compared to Col-0 largely differed among the accessions from south-west of France, with values ranging from 0.2 to more than 1.5 (Supplementary Fig. S1). In comparison, this ratio for the non-adherent mucilage shows less variability among populations, with values ranging from 0.5 to 1.5 (Supplementary Fig. S1). Note that the sizes of both mucilage layers are positively correlated with seed size (Supplementary Table S1).

On average, a large and significant genetic variation was detected among accessions within populations (Table 1), with the observation of populations presenting either no phenotypic or extensive variation among accessions for adherent or non-adherent mucilage areas (Fig. 2). Overall, while 32.3 % and 13.8 % of phenotypic variance for adherent and non-adherent mucilage areas result from a genetic differentiation among populations, respectively, a higher proportion of phenotypic variance result from a genetic differentiation among accessions within populations, i.e. 42.7% and 34.1% for adherent or non-adherent mucilage areas, respectively (Table 1).

**Fig. 2.**
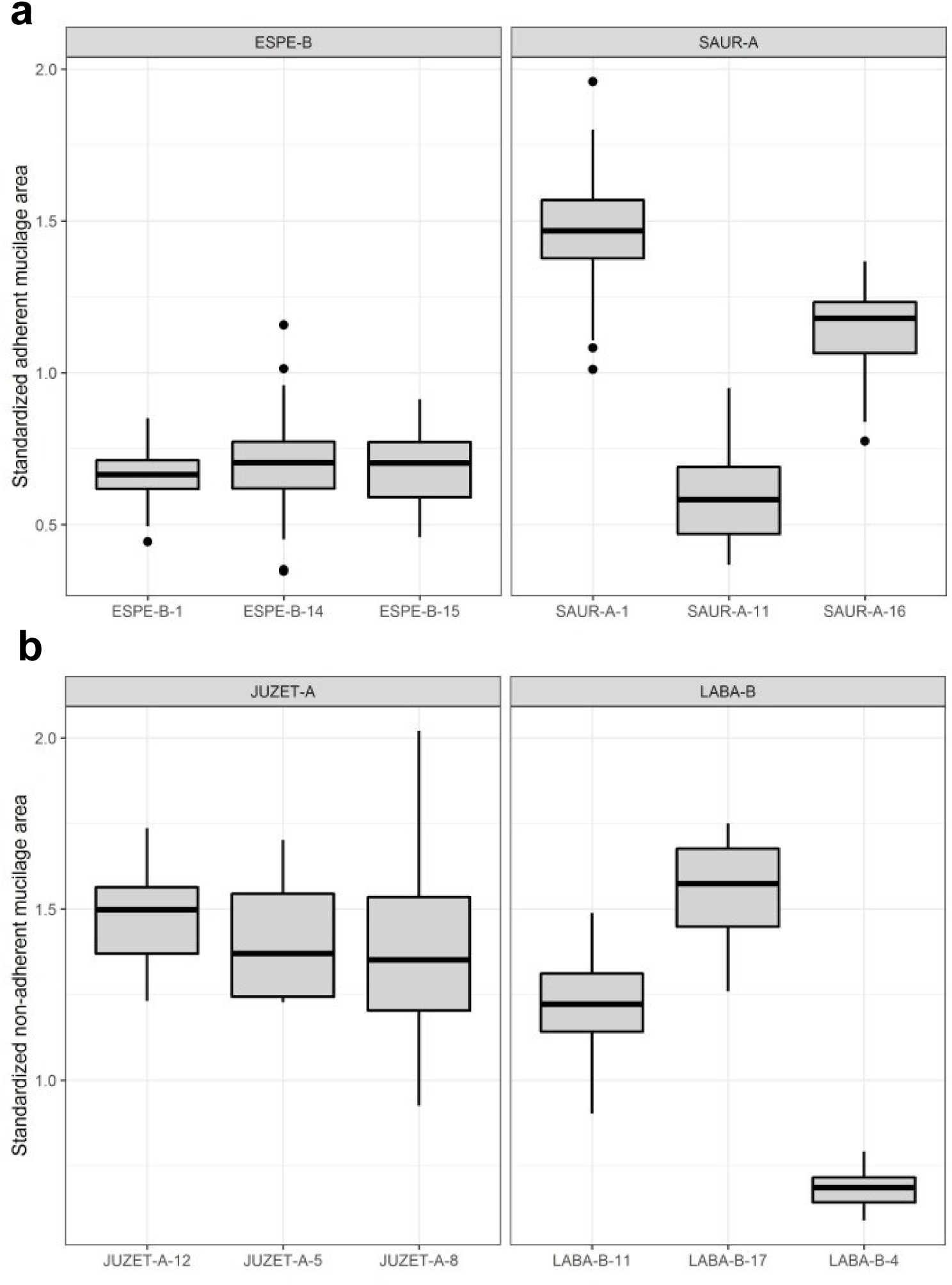
The populations can contain either homogeneous or contrasted accessions for each mucilage layers. Example of adherent **(a)** and non-adherent **(b)** mucilage standardized area values for four population examples containing three homogeneous (left side) and three contrasted (right side) accessions, respectively.

**Table 1.** Global statistics for the used GWAS model shows that variance is surprisingly more explained by morphological variability between the 424 accessions than between the 166 populations. *F* stands for a F-test, LTR for likelihood ratio test, *P* for the P-value and *R*^2^ for coefficient of determination ("R squared")

| Terms | Adherent mucilage area |  |  | Non adherent mucilage area |  |  |
| --- | --- | --- | --- | --- | --- | --- |
|  | *F* or LRT | *P* | *R*<sup>2</sup> | *F* or LRT | *P* | *R*<sup>2</sup> |
| Population | 2,9 | 3,2E-14 | 32,3 | 1,82 | 1,1E-05 | 13,8 |
| Accession (Population) | 30514,8 | 1,0E-16 | 42,7 | 642,3 | 1,0E-16 | 34,1 |
| Seed area | 13811,8 | 1,0E-32 | - | ne | ne | ne |
| Residuals |  |  | 25,0 |  |  | 52,1 |

No correlation was observed at the population level between the standardized adherent mucilage area and standardized non-adherent mucilage area (Spearman’s *rho* 0.09, *P =* 0.27) (Fig. 3).

**Fig. 3.**
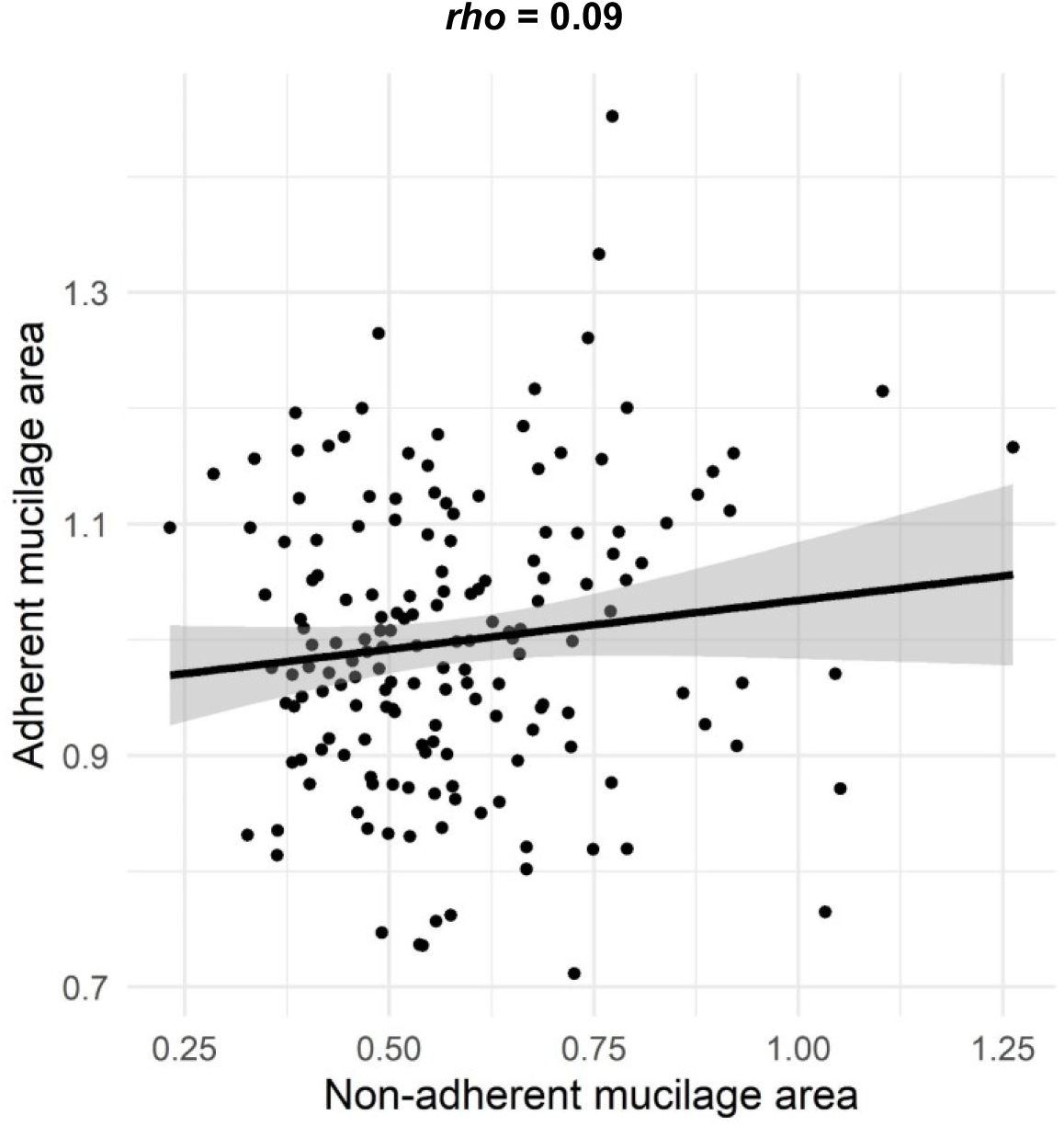
The two mucilage areas are not correlated at the population level. Scatter plots of the mean value of each population for the standardized adherent mucilage versus standardized non-adherent mucilage areas. The black line corresponds to the linear model with its confidence interval in grey. Each dot corresponds to one population containing data from its accessions. *rho* is the spearman correlation coefficient.

### 3.2 Relationship between mucilage and environmental abiotic parameters

Among population, variation of the standardized adherent mucilage area was significantly correlated with several abiotic parameters collected from their native sites (Frachon et al., 2019, 2018) (Fig. 4). The area of the adherent mucilage was negatively correlated with the mean of annual temperatures (Fig. 4a) and the mean of coldest month temperatures (Fig. 4b) and positively correlated with the precipitation level during winter (Fig. 4c) and autumn (Fig. 4d). In contrast, variation of the non-adherent mucilage area among populations was not correlated with any climate variables (Table 2, Supplementary Table S1). In addition, while the adherent seed mucilage area was weakly associated with the total concentration in nitrogen (positive relationship) and pH (negative relationship) (Supplementary Table S1), no significant correlation was detected for the non-adherent mucilage with any of the variables describing the agronomic properties of the soil of the native sites (Table 2, Supplementary Table S1).

**Fig. 4.**
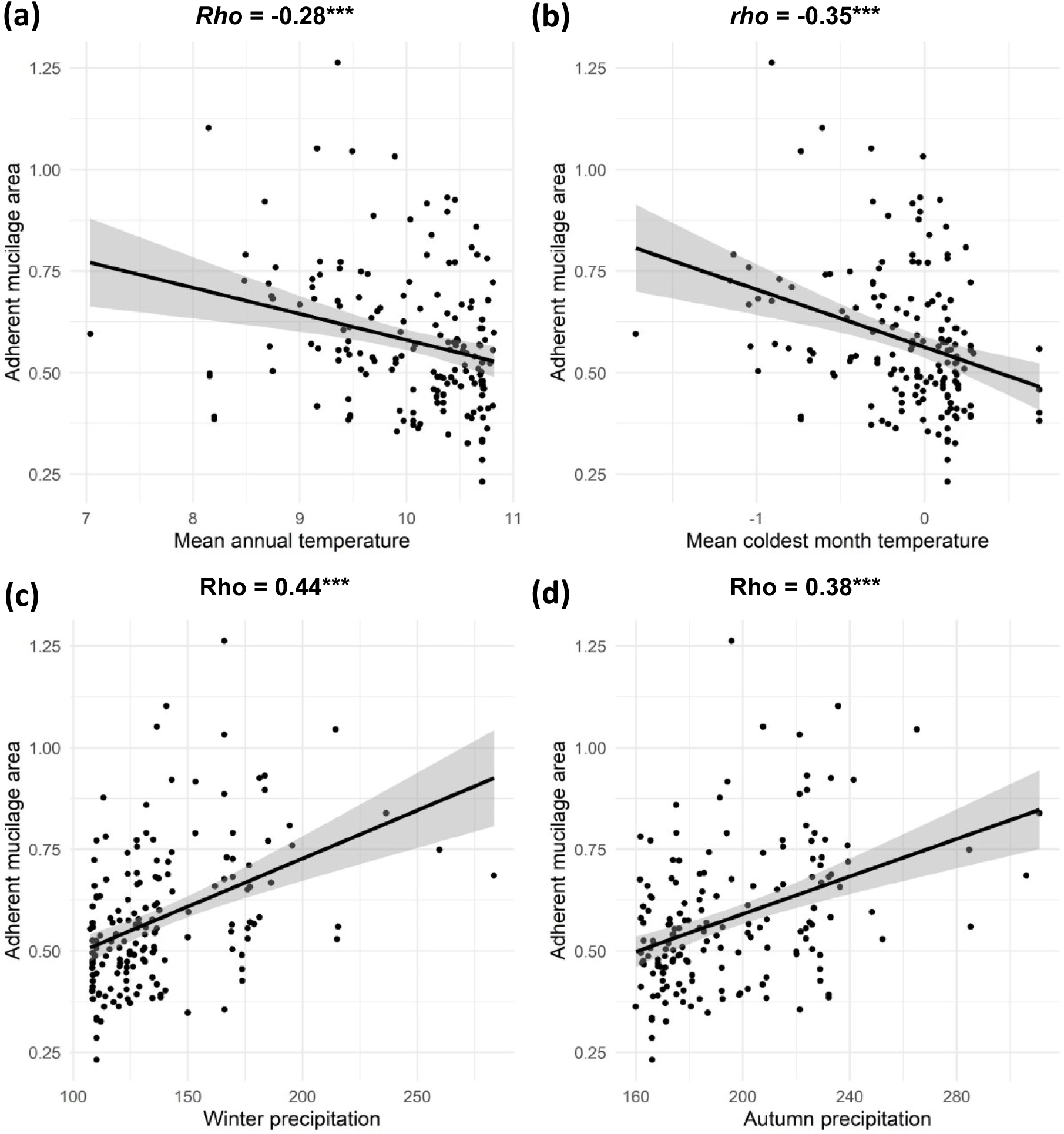
The adherent mucilage area is negatively correlated with temperature and positively with precipitation. Graphs show the relationship between the standardized adherent mucilage area and the mean annual temperature **(a)**, mean coldest month temperature **(b)**, winter precipitation **(c)** and autumn precipitation **(d)**. The Spearman correlation coefficient is indicated above each graph. The black line is the linear model with its confidence interval in grey. These four environmental parameters are the only ones to remain significant (P-value < 0.001) after FDR correction of P-values among the tested parameters (Supplementary Table S1). Each dot corresponds to one population containing data from its accessions.

**Table 2.**
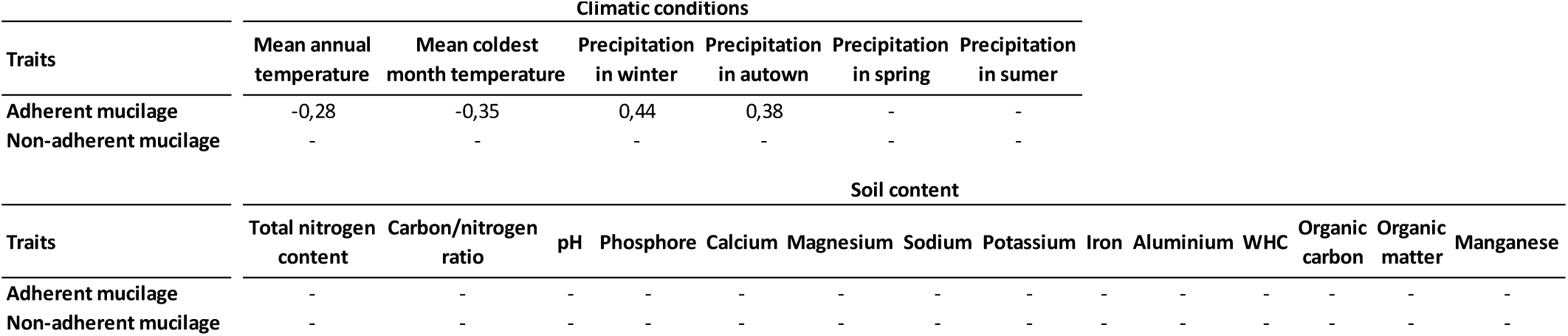
Spearman correlation across every populations between measured traits and their harvest site environmental parameters. Indicated values are correlation coefficients that are significant after FDR correction.

### 3.3 The phenotypic variation of the standardized areas of seed mucilage layers is highly polygenic

GWA analyses revealed a polygenic architecture for both phenotypic traits, with the detection of 26 QTLs overlapping with 55 genes associated with the adherent mucilage area variability and 10 QTLs overlapping with 28 genes associated with the non-adherent mucilage area variability (Fig. 5, Supplementary Fig. S2 and S3). The 36 QTL(corresponding to 83 ORFs) encompass 512 SNPs distributed as following 198 in the coding regions (1 stop gained, 71 non synonymous mutations and 126 synonymous mutations), 177 in the non coding regions and 137 upstream or downstream of the genes (Supplementary Table S4). Among these 36 QTLs, none were simultaneously associated with both mucilage layers.

**Fig. 5.**
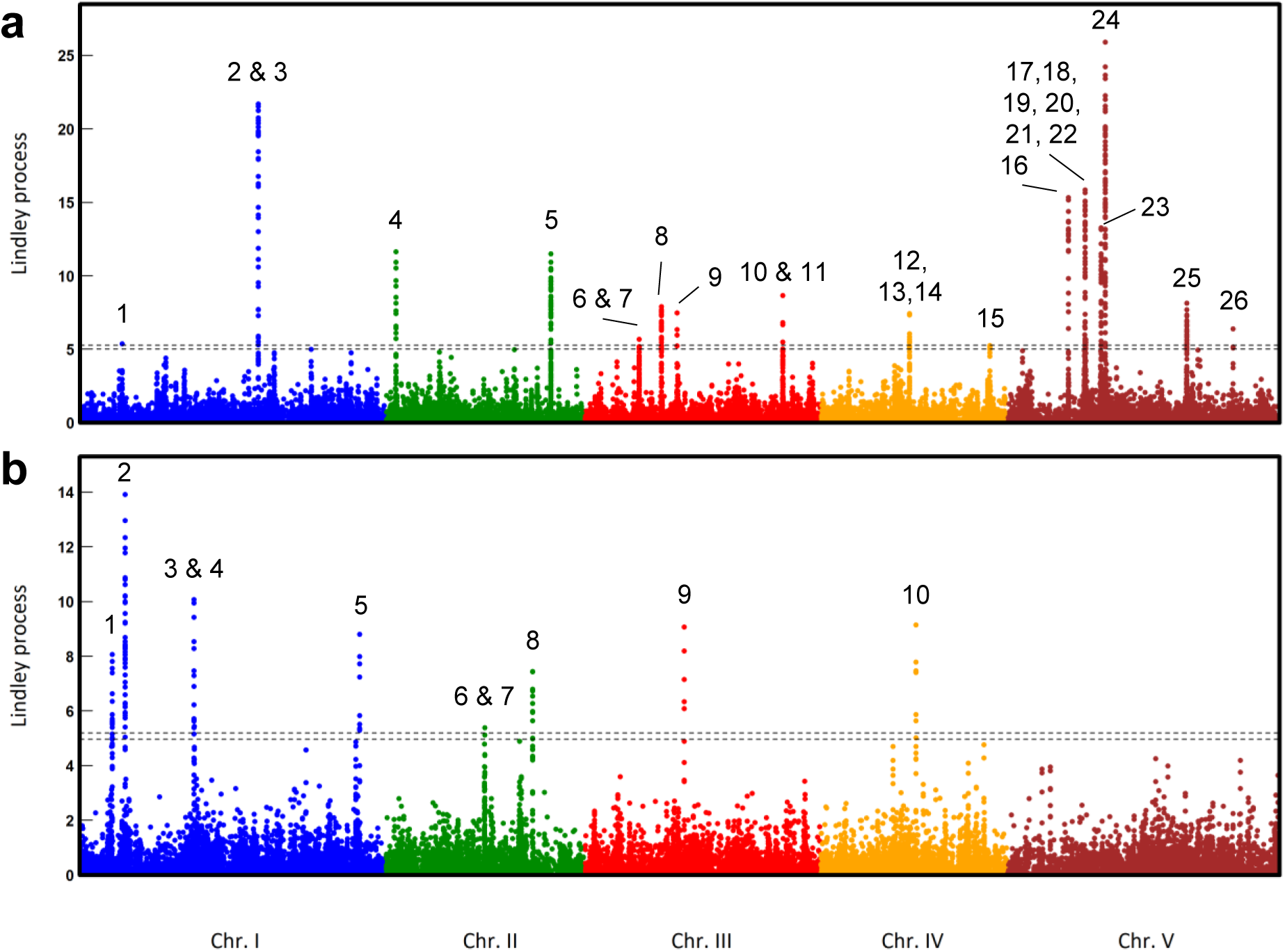
Highlight of genome QTLs explaining the morphological variability of adherent and non-adherent seed mucilage standardized area. Manhattan plots representing the Lindey process value for each SNP among the genome of *A. thaliana*, highlights 26 QTLs for adherent **(a)** and 10 QTLs for non-adherent **(b)** seed mucilage, respectively. The threshold used for QTL selection is represented by the dotted line at Lindey value of 5. Everything below is considered as background noise.

Some of the 83 genes identified in our study are members of multigenic families containing already described myxospermy-related genes. For instance, we identified *GA2OX4* (*At1g47990*) and *SK19* (*At2g03160*) for adherent mucilage (Fig. 6a), and *HB-2* (*At4g16780*), *SK41* (*At1g09840*), *CuAOα2* (*At1g31690*), and *CuAOα3* (*At1g31710*) for non-adherent mucilage (Fig. 6b). These candidate genes belong to four independent multigenic families containing gene members of the seed mucilage toolbox: *GA3OX4* (*At1g80330;* (Kim et al., 2005)), *SK11*/*SK12* (*At4g34210*/*At4g34470*; (Li et al., 2018)), *HB-25* (*At5g65410*; (Bueso et al., 2014)), and *CuAOα1* (*At1g31670*, (Fabrissin et al., 2019)). Interestingly, two multigenic families not previously shown to be involved in myxospermy were overrepresented, such as the cytochrome P450 (CYP450) family with six genes (*At2g34500*, *At4g15380*, *At3g25180*, *At4g15393*, *At2g34490*, *At1g13110*) and the serine carboxypeptidase like (SCPL) family with 4 genes (*At2g23000*, *At2g22980*, *At2g22970*, *At2g22990*) (Fig. 6).

**Fig. 6.**
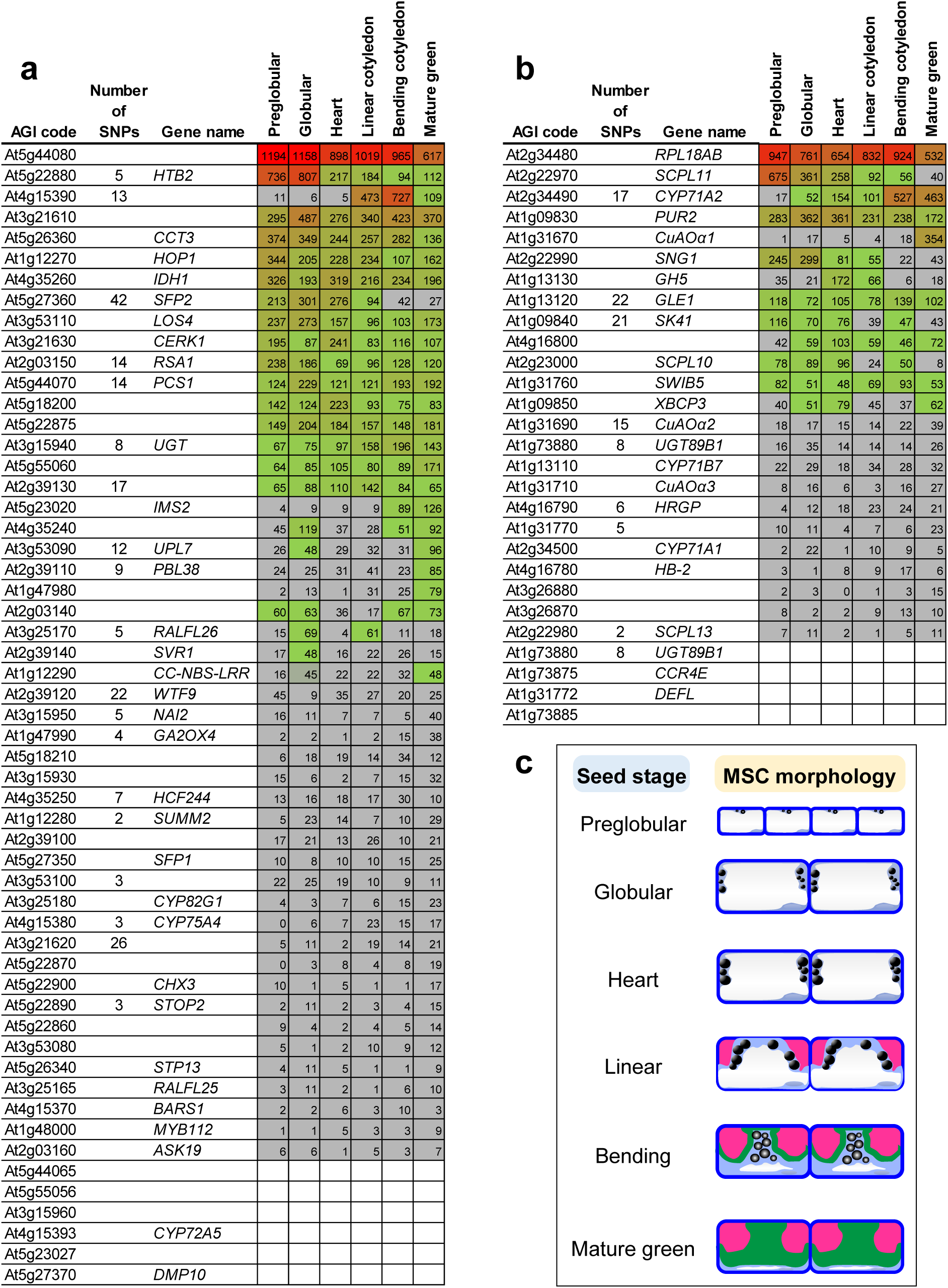
About 50% of the 83 candidate genes from the GWAS are expressed in the seed coat during seed development. Fifty five and 28 selected genes are present within 26 and 10 QTLs highlighted by the GWAS for adherent **(a)** and non-adherent **(b)** mucilage layers, respectively. These are confronted to previously published transcriptomic data of the laser-captured seed coat during six seed developmental stages (Belmonte et al., 2013). No significant values below the threshold of “45” are in grey, low expressed genes are in green, highly expressed genes are in red. Note that ten genes were not present in the tissue arrays. Number of significant SNPs found within the gene region are indicated, genes with no value were located downstream or upstream of significant SNPs. **(c)** Mucilage secretory cell (MSC) morphology for every seed developmental stage available in transcriptomic (adapted from Francoz et al., 2019).

*SFP2* (*At5g27360*) is a candidate gene identified within the most significant QTL associated with the adherent mucilage area (i.e. QTL 24, Fig. 5) with the presence of 42 significant SNPs within the gene (Supplementary Table S3). Interestingly, *SFP2* is expressed during early seed development (Fig. 6a). Worth of noting, QTL 24 also contains *SFP1* (*At5g27350*), the tandem duplicated of *SFP2*, but notexpressed during seed development (Fig. 6a). For the non-adherent mucilage area, an interesting polysaccharide-related candidate gene is *At1g13130*, an uncharacterized glycosyl hydrolase family 5 (GH5) found upstream of 22 significant SNPs within QTL 2 (Supplementary Table S3).

## 4 Discussion

In this study, we aimed at (i) exploring the natural genetic variation in myxospermy among 166 natural populations of *A. thaliana* collected at a regional scale, (ii) identifying the ecological factors acting as putative drivers of myxospermy, and (iii) applying recent methodologies in GWA mapping to unravel the underlying genetic architecture.

The whole study is based on seed mucilage area variation. The main concern was the slight positive correlation of seed size with the size of both mucilage layers, (Supplementary Table S1). If the cells keep a proportional size in the seed, it is coherent to have larger mucilage secretory cells in larger seeds (not evaluated here) and, consequently, more abundant mucilage. Nonetheless, extensive genetic variation was detected for both mucilage traits after correcting for the effect of seed size variation, using seed size as a covariate (Table 1).

Adherent and non-adherent mucilage have different polysaccharide compositions that provide structure only to the former (Macquet et al., 207b). According to the hypothesis that for an equal amount of synthetized mucilage, the adherent and non-adherent mucilage areas are dependent on their structural polysaccharide content, the two layers of mucilage should be negatively correlated. However, adherent mucilage and non-adherent mucilage were uncorrelated in this study (Fig. 3), thereby contradicting this hypothesis. Consequently, the differences of mucilage area among accessions could be due to differences in total amount of synthetized mucilage and/or in a change in mucilage composition. Then, in this scenario, we have more likely measured a value that depends on a combination of mucilage amount and composition. Note that the experimental estimation of the area of the non-adherent mucilage could also reflect its ability to spread on agarose, then its structure, rather than its total amount.

Variation among populations in adherent mucilage is negatively correlated with temperature. It was recently demonstrated that the thick adherent seed mucilage of *Lepidium perfoliatum*, another myxospermous Brassicaceae species, prevents the germination under relatively low temperatures (10°C and 15°C) and increases germination under higher temperatures (25°C and 30°C), in particular in presence of a long light time exposure (Zhou et al., 2021). This suggests a role of the adherent mucilage in germination inhibition under cold temperature until optimal conditions are obtained. The observed correlation between adherent mucilage and temperature and precipitation could therefore be indirect, through germination regulation. Indeed, *A. thaliana* seed mucilage was often assumed to favor germination by providing more water to the embryo (Arsovski et al., 2009; Penfield, 2001). However, 24 h after imbibition, *A. thaliana* embryos contain less water in mucilaginous seeds than in non-mucilaginous seeds (Saez-Aguayo et al., 2014). In *L. perfoliatum*, seed mucilage was shown to increase drastically germination in optimal water content whereas no impact was found under osmotic stress (Zhou et al., 2021). The absence of correlation between soil composition and non-adherent mucilage might be explained by the variability of soil composition at the micro-scale level and also because the non-adherent mucilage will rapidly spread into the ground during the first rain and thus could not influence directly the seed development. However, as the non-adherent mucilage spreads into the ground, it is surprising that physical parameters such as soil water holding capacity are not correlated. Therefore, the fact that mucilage phenotypes are rather related to temperature and precipitation than soil properties suggests a possible physiological importance of seed mucilage during underground seedling development rather than during germination.

Fewer QTLs were found for the non-adherent mucilage than for the adherent mucilage, which is in agreement with the estimates of inter-population variance (13.8 % for non-adherent mucilage compared to 32.3 % for adherent mucilage). In *A. thaliana*, about one hundred genes have previously been characterized for their implication in seed mucilage biosynthesis and release (Viudes et al., 2021). Our GWA mapping approach allowed identifying 82 new candidate genes and only one belonging to the list of previously validated candidate genes. The 100 genes already characterized from previous studies together with our GWA analyses highlight the extreme polygenic character for mucilage traits. Our GWAS revealed genes inducing weak phenotypes more difficult to identify with regular genetic approach (reverse and forward genetics), which can in part explain the small overlap between the two sets of identified genes.

The gene already known for its implication in seed mucilage is *CuAOα1* (*At1g31670*) encoding a copper amine oxidase that was identified with GWAS on non-adherent mucilage (Supplementary Fig. S3 and Table S3). The *cuaoα1* mutant has less rhamnogalacturonan I (RGI) pectin domain in its non-adherent mucilage (Fabrissin et al., 2019). Interestingly, *CuAOα1* has been identified with two other genes in a previous GWAS on mucilage, which did not reveal any other previously characterized myxospermy-related genes (Fabrissin et al., 2019; Voiniciuc et al., 2016) illustrating the pionner capacity of this approach. In addition to its expression in seeds that is restricted to the seed coat, *CuAOα1* is expressed in root tips (Klepikova et al., 2016), especially in the lateral root cap and it is induced by high nitrate concentration (Gifford et al., 2008). During the lateral root cap cell formation, a pectinaceous mucilage accumulates inside the sixth cell layer (c6) below the quiescent center to, *in fine* extrude root mucilage around the c7 (Maeda et al., 2019). Because CuAOα1 seems to be involved in RGI production in seed mucilage (Fabrissin et al., 2019), it would be interesting to investigate as well its putative role in root mucilage. Although a putative link between root mucilage and non-adherent mucilage remains elusive, there is an interesting parallel between their developmental and ecological roles.

The identification of new putative candidate genes is also well supported by their spatio-temporal expression. Among the 83 candidates, 40 are expressed in the seed coat along the Col-0 seed development kinetics (Belmonte et al., 2013) (Fig. 6). Previously characterized functions, associated with seed development expression data, provide interesting elements for some candidates. Intriguingly, several candidate genes are related to the secondary metabolism. *IMS2*/*MAM3* (*At5g23020*), with a role in glucosinolate biosynthesis (Petersen et al., 2019), is located upstream of 10 significant SNPs within the adherent mucilage QTL 21 (Supplementary Table S3) and expressed only at the mature green stage, just before seed drying (Fig. 5 and Fig. 6a). The *CYP450* gene, *CYP71A2* (*At2g34490*), contains the 17 significant SNPs of the non-adherent mucilage QTL 8 and shows a seed coat-specific expression increasing along seed development (Fig. 6b) and extending to endosperm at mature green stage (Supplementary Table S2). *CYP450* genes can be implicated in seed suberin and cutin biosynthesis (Renard et al., 2020) and are often implicated in secondary metabolite decoration (Schuler and Werck-Reichhart, 2003). *SCPL11* (*At2g22970*), *SNG1* (*At2g22990*), and *SCPL10* (*At2g23000*), localized within non-adherent QTLs 6 and 7 (Fig. 5), have a seed coat specific expression. These three genes are serine carboxypeptidase-like proteins that belong to a large multigenic family implicated in secondary metabolism. *SCPL19*/*SNG2* (*At5g09640*) has a role in sinapoylcholine formation in *A. thaliana* seeds (Shirley et al., 2001). Although a direct link between *A. thaliana* seed mucilage and secondary metabolism remains elusive, a regulatory network was identified between seed mucilage, seed pigments and seed secondary metabolites (Lepiniec et al., 2006; Salem et al., 2017; Viudes et al., 2020). *SFP1*, the paralog of *SFP2* that is located within QTL 24, encodes a monosaccharide transporter during leaves senescence (Quirino et al., 2001). The involvement of SFP2 in sugar transport related to seed adherent mucilage polysaccharides remains to be demonstrated. *At1g13130* found upstream of 22 significant SNPs is a GH5 whose substrate specificity is unknown. However, GH5 family members have glycosyl hydrolase activities on various substrates (http://www.cazy.org/GH5.html) including polysaccharides found in the mucilage secretory cells (MSCs) such as cellulose, xylan or mannan and might be implicated in mucilage polysaccharide hydrolysis. *At1g13130* is expressed sharply at heart and linear cotyledon stages (Fig. 6a), co-occurring partially with cellulose fibrils formation since *CESA5* (*At5g09870*) (main cellulose synthase involved in seed mucilage) is highly expressed at the linear cotyledon stage (Sullivan et al., 2011).

Additionally, in other species than *A. thaliana*, seed and root mucilages were shown to change soil physical properties, which in turn influence water retention and water availability (Ahmed et al., 2016; Deng et al., 2014), and induce modification of the rhizospheric microbial communities (Sasse et al., 2018). If the seed mucilage is also implicated in biotic interactions, it may explain the association of both seed mucilage layers with several candidate genes implicated in secondary metabolism identified in our GWAS, knowing that plant secondary metabolites play crucial roles during plant-microbes interactions (Kessler and Kalske, 2018).

## Conclusion

The seed mucilage was highly variable among the Midi-Pyrénées regional populations for both layers. The detected QTLs underlying this morphological variability, contain one previously characterized gene and 82 putative new seed mucilage actors. Considering the 100 genes already characterized in previous studies, this highlights the extreme polygenic character of this complex trait in *A. thaliana*. The candidate gene *CuAOα1* (*At1g31670*) identified in our study for the non-adherent mucilage, was previously shown to be involved in non-adherent mucilage, thereby validating the significance of our approach. New candidate genes coming from our study await functional validation. The correlation of adherent seed mucilage area with temperature and precipitations seems to be important and the tempting indirect link of this correlation with germination will be interesting to validate through germination assays using seed with or without mucilage. Altogether this study shed light on putative new genetic actors and on the ecological role of seed mucilage.

## Acknowledgements

The authors are thankful to the Paul Sabatier-Toulouse 3 University and to the Centre National de la Recherche Scientifique (CNRS) for granting their work. This work was also supported by a grant by The Fédération de Recherche Agroscience Interactions Biodiversité (https://www.fraib.fr/fraib_eng/). We thank Werner Schweyer for his technical support during seed phenotyping.

---

**Supplementary Fig. S1.**
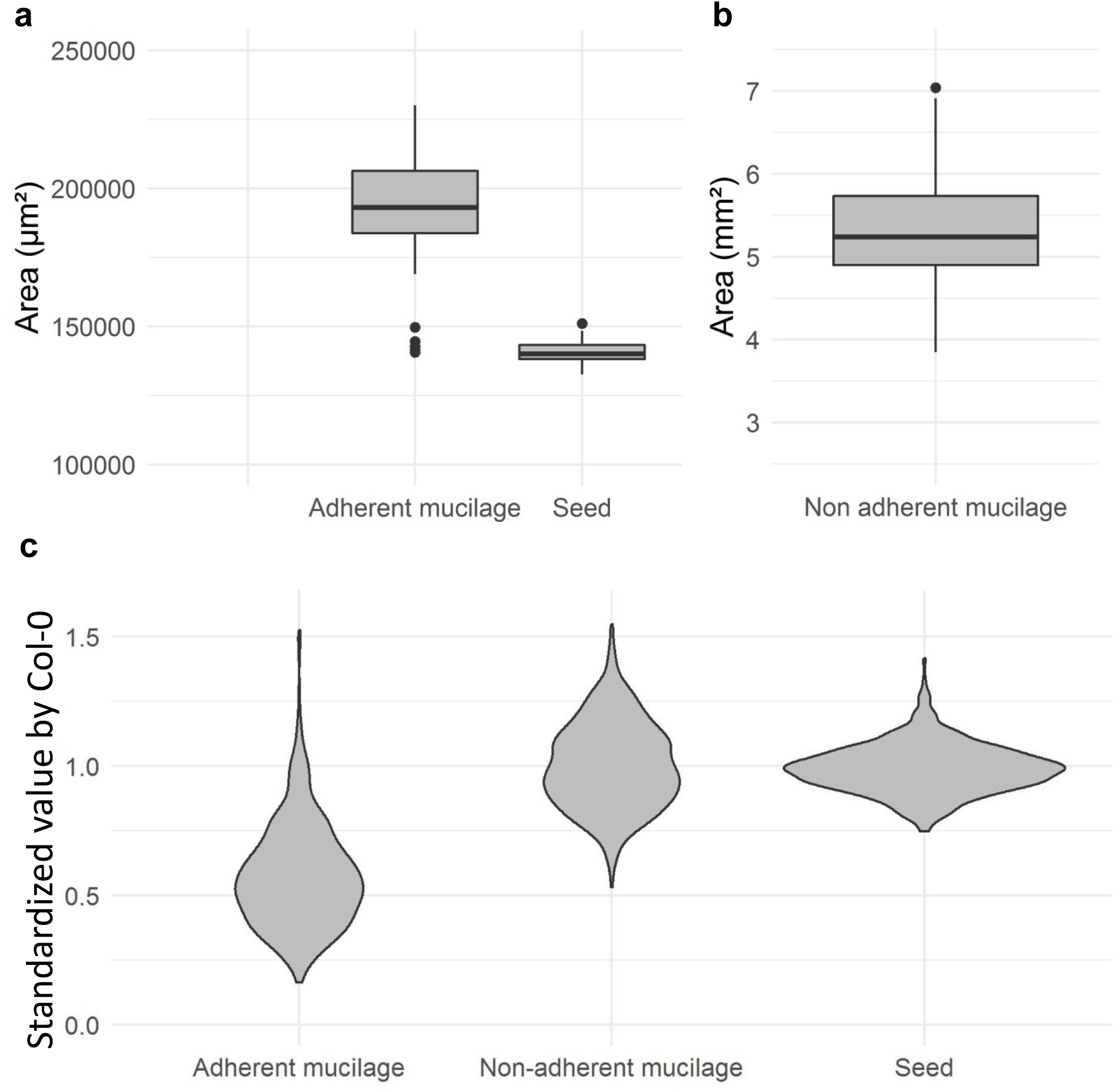
Relatively low variation of the three parameters measured on the Col-0 standard included in each batch and repartition of the observed variability of the studied natural populations phenotypes after normalization with the internal Col-0 standard. **(a)** Boxplot showing mean values of seed area and adherent mucilage area of the Col-0 seeds used as a standard in the 44 individual experimental batches during the phenotyping of studied populations. The areas are expressed in µm². (**b)** Boxplot showing mean values of non-adherent mucilage areas of the Col-0 seeds used as a standard in the 98 batches of experiment during the phenotyping of studied populations. The areas are expressed in mm². (**c)** Violin plot showing mean values of seed, adherent mucilage, and non-adherent mucilage area normalized with the Col-standard (y axis) for each of the 424 accessions. Area values are expressed relatively to the mean area of Col-0 seeds (set to 1) included in each respective experimental batch. The x axis wideness of violin plot shows the relative frequency of accessions for the corresponding value displayed on the y axis.

**Supplementary Fig. S2.**
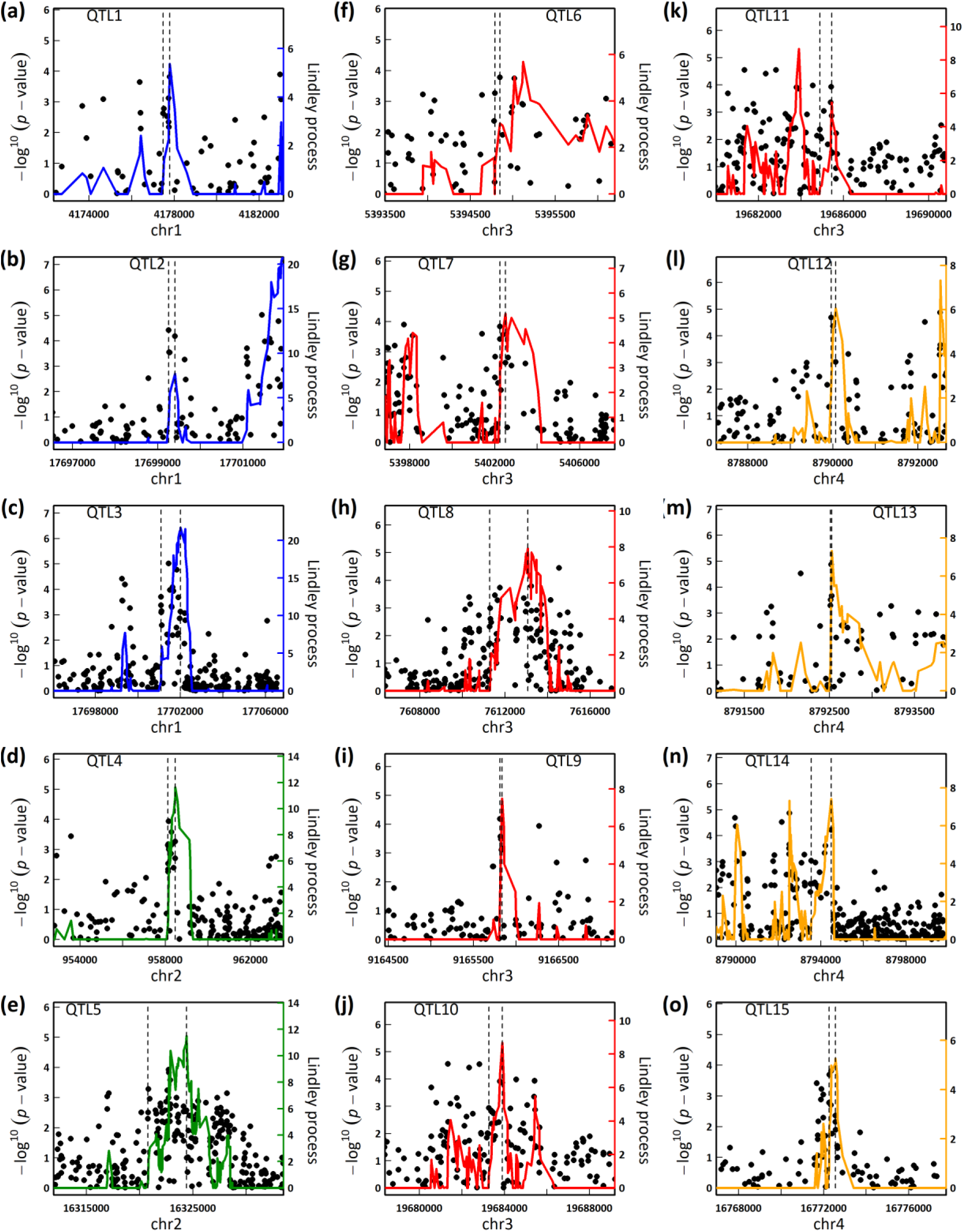
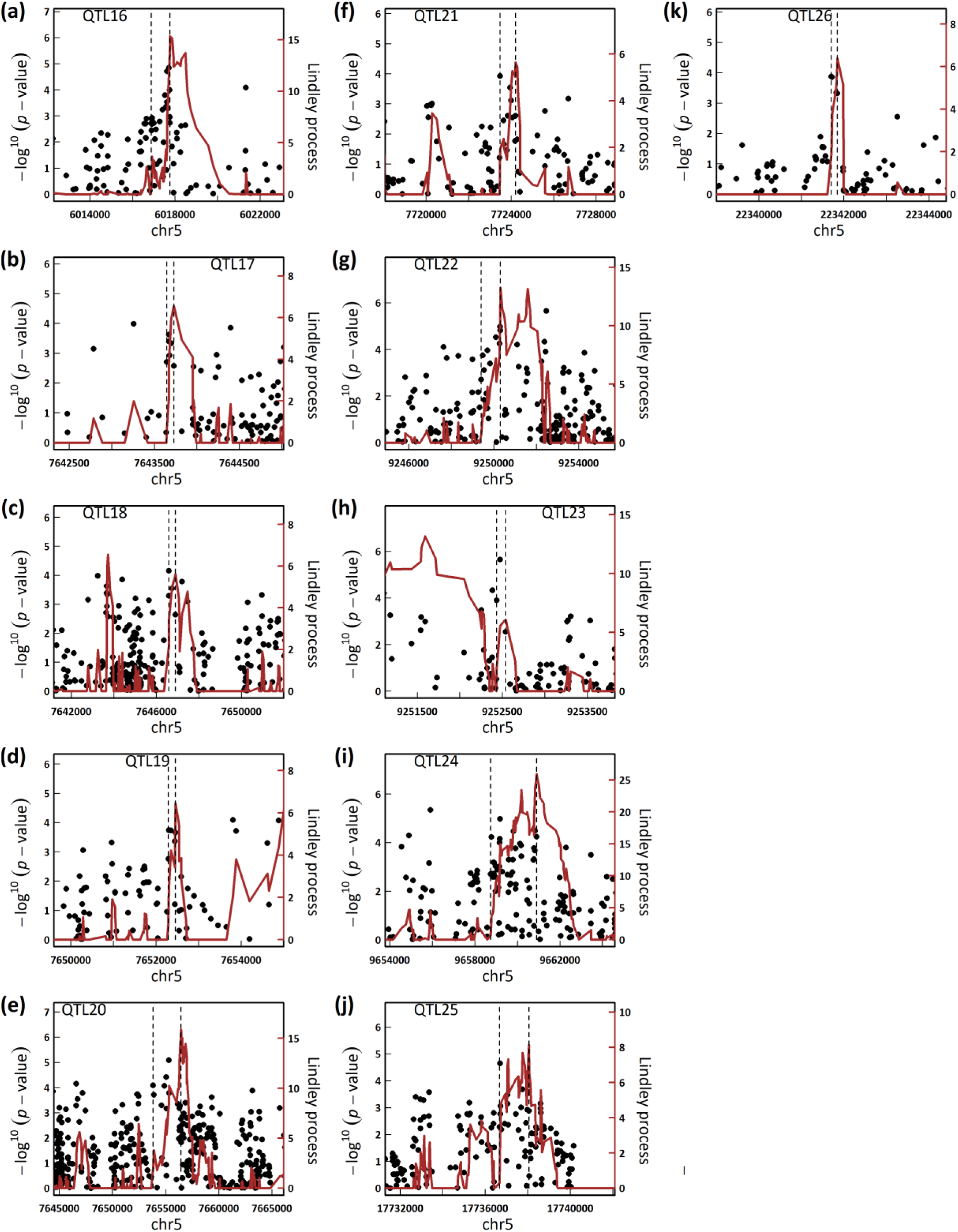
Zoom on significant QTLs related to the genetic variability of adherent mucilage. Each peak is delimited by the two vertical dot lines with the first and last significant SNP contained in it. Each SNP is ploted with its –log10 of the P-value and the colored line represents the Lindley process evolution.

**Supplementary Fig. S3.**
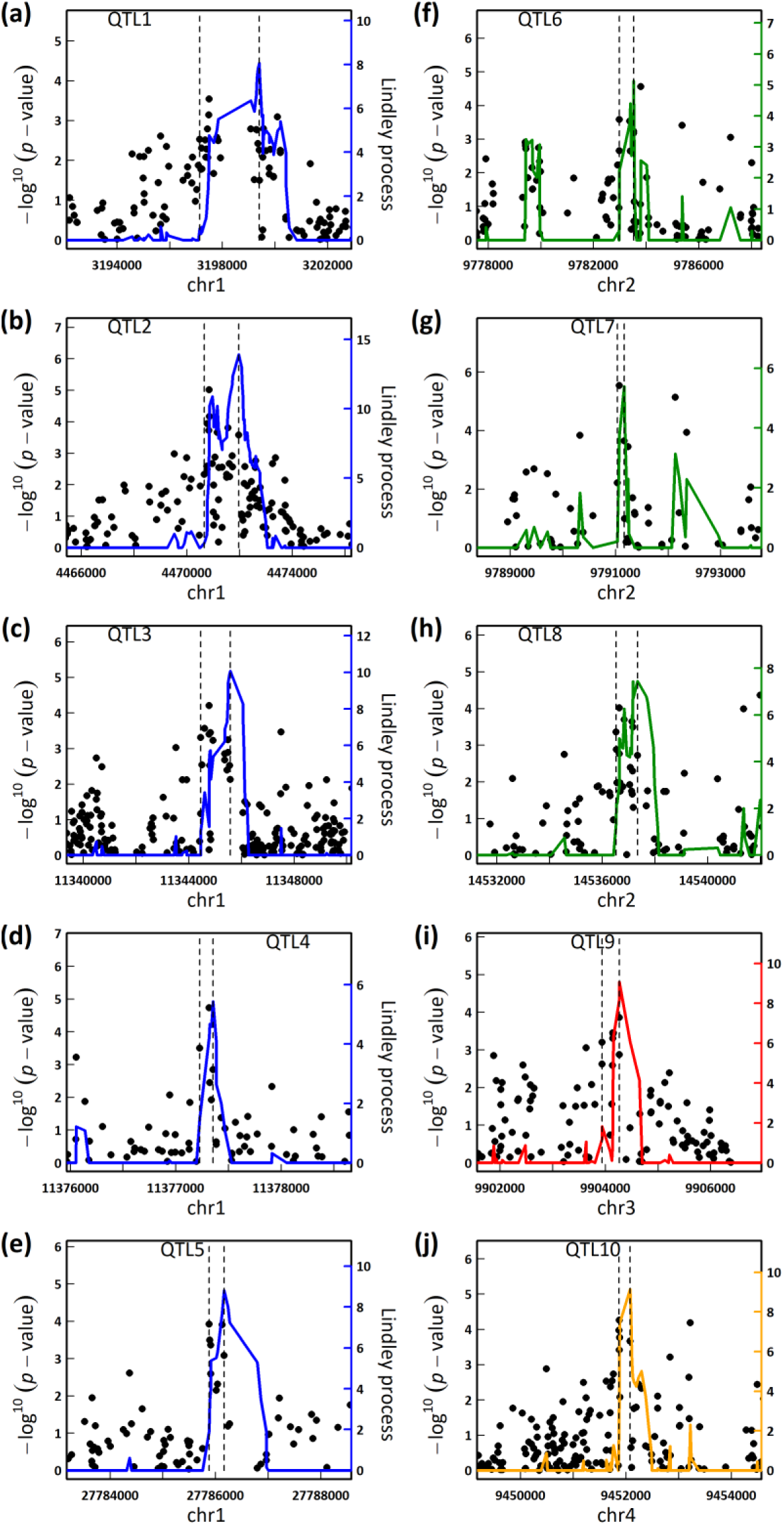
Zoom on significant QTLs related to the genetic variability of non-adherent mucilage. Each peak is delimited by the two vertical dot lines with the first and last significant SNP contained in it. Each SNP is ploted with its –log10 of the P-value and the colored line represents the Lindley process evolution.

**Supplementary Table S1.** Spearman correlation matrix across every population between measured traits and their harvested site environmental parameters. Indicated values in the upper half are spearman correlation coefficient and values in the down part are the coresponding p-values after an FDR correction.

**Supplementary Table S2.** Expression of every candidate genes in the different seed tissues during six seed developmental stages (Belmonte et al., 2013). No significant values below the threshold of “45” are in grey, low expressed genes are in green, highly expressed genes are in red. Note that 10 genes had no spot ID in the tissue arrays.

**Supplementary Table S3.** Distribution and number of significant SNPs in each QTLs. The Table shows the number of SNPs identified with the GWAS performed on the adherent mucilage (left) and non-adherent mucilage (right) and ID of the gene overlapping with each SNP, or found within a 2 kb upstream and downstream region.

| Adherent mucilage |  |  |  |  | Non-adherent mucilage |  |  |  |  |
| --- | --- | --- | --- | --- | --- | --- | --- | --- | --- |
| QTLs | SNPs | Overlap ID | Upstream ID | Downstream ID | QTLs | SNPs | Overlap ID | Upstream ID | Downstream ID |
| QTL1 | 2 | AT1G12280 | AT1G12270 | AT1G12290 | QTL1 | 21 | AT1G09840 | AT1G09830 | AT1G09850 |
|  | 5 | NA | AT1G12280 | AT1G12290 | QTL2 | 22 | AT1G13120 | AT1G13110 | AT1G13130 |
| QTL2 | 4 | AT1G47990 | AT1G47980 | AT1G48000 | QTL3 | 15 | AT1G31690 | AT1G31670 | AT1G31710 |
| QTL3 | 23 | NA | AT1G47990 | AT1G48000 | QTL4 | 5 | AT1G31770 | AT1G31760 | AT1G31772 |
| QTL4 | 14 | AT2G03150 | AT2G03140 | AT2G03160 | QTL5 | 8 | AT1G73880 | AT1G73875 | AT1G73885 |
| QTL5 | 9 | AT2G39110 | AT2G39100 | AT2G39120 | QTL6 | 2 | AT2G22980 | AT2G22970 | AT2G22990 |
|  | 1 | NA | AT2G39110 | AT2G39120 |  | 4 | NA | AT2G22980 | AT2G22990 |
|  | 22 | AT2G39120 | AT2G39110 | AT2G39130 | QTL7 | 3 | NA | AT2G22990 | AT2G23000 |
|  | 17 | AT2G39130 | AT2G39120 | AT2G39140 | QTL8 | 17 | AT2G34490 | AT2G34480 | AT2G34500 |
| QTL6 | 8 | AT3G15940 | AT3G15930 | AT3G15950 | QTL9 | 10 | NA | AT3G26870 | AT3G26880 |
| QTL7 | 5 | AT3G15950 | AT3G15940 | AT3G15960 | QTL10 | 6 | AT4G16790 | AT4G16780 | AT4G16800 |
| QTL8 | 26 | AT3G21620 | AT3G21610 | AT3G21630 |  |  |  |  |  |
| QTL9 | 5 | AT3G25170 | AT3G25165 | AT3G25180 |  |  |  |  |  |
|  | 12 | AT3G53090 | AT3G53080 | AT3G53100 |  |  |  |  |  |
| QTL11 | 5 | AT3G53090 | AT3G53080 | AT3G53110 |  |  |  |  |  |
|  | 3 | AT3G53100 | AT3G53090 | AT3G53110 |  |  |  |  |  |
| QTL12 | 3 | AT4G15380 | AT4G15370 | AT4G15390 |  |  |  |  |  |
| QTL13 | 4 | NA | AT4G15380 | AT4G15390 |  |  |  |  |  |
| QTL14 | 13 | AT4G15390 | AT4G15380 | AT4G15393 |  |  |  |  |  |
|  | 1 | NA | AT4G15390 | AT4G15393 |  |  |  |  |  |
| QTL15 | 7 | AT4G35250 | AT4G35240 | AT4G35260 |  |  |  |  |  |
| QTL16 | 20 | NA | AT5G18200 | AT5G18210 |  |  |  |  |  |
| QTL17 | 6 | NA | AT5G22860 | AT5G22870 |  |  |  |  |  |
| QTL18 | 4 | NA | AT5G22860 | AT5G22870 |  |  |  |  |  |
| QTL19 | 5 | AT5G22880 | AT5G22875 | AT5G22890 |  |  |  |  |  |
| QTL20 | 3 | AT5G22890 | AT5G22880 | AT5G22900 |  |  |  |  |  |
|  | 16 | NA | AT5G22890 | AT5G22900 |  |  |  |  |  |
| QTL21 | 10 | NA | AT5G23020 | AT5G23027 |  |  |  |  |  |
| QTL22 | 14 | NA | AT5G26340 | AT5G26360 |  |  |  |  |  |
| QTL23 | 3 | NA | AT5G26340 | AT5G26360 |  |  |  |  |  |
| QTL24 | 42 | AT5G27360 | AT5G27350 | AT5G27370 |  |  |  |  |  |
| QTL25 | 14 | AT5G44070 | AT5G44065 | AT5G44080 |  |  |  |  |  |
|  | 5 | NA | AT5G44070 | AT5G44080 |  |  |  |  |  |
| QTL26 | 4 | NA | AT5G55056 | AT5G55060 |  |  |  |  |  |

